# Assessment of Human Expertise in First-Person Shooter Games

**DOI:** 10.1101/2022.06.30.498231

**Authors:** Ian Donovan, Marcia A. Saul, Kevin DeSimone, Jennifer B. Listman, Wayne E. Mackey, David J. Heeger

## Abstract

Contrary to traditional professional sports, there are few standardized metrics in professional esports (competitive multiplayer video games) for assessing a player’s skill and ability. To begin to address this, we assessed the performance of professional-level players in Aim Lab™, a first-person shooter training and assessment game, within two separate target-shooting tasks. These tasks differed primarily in the relative incentive for fast and imprecise shots versus slow and precise shots. Each player’s motor acuity was measured by characterizing the speed-accuracy trade-off in shot behavior: shot frequency and shot spatial error (distance from center of a target). We also characterized the fine-grained kinematics of players’ mouse movements. Our findings demonstrate that: 1) movement kinematics depended on task demands; and 2) individual differences in motor acuity were significantly correlated with both kinematics and the number of movements needed to hit a target. We demonstrate the importance of transforming from orientation in the virtual environment to centimeters on the mouse pad, as well as accounting for differences in mouse sensitivity across players, for characterizing human performance in first-person shooter games. This approach to measuring motor acuity has widespread application not only in esports assessment and training, but also in basic (motor psychophysics) and clinical (gamified rehabilitation) research.

## 1 INTRODUCTION

The principled study of digital game performance is in its infancy (Campbell et al., 2018; Huang et al., 2017), even though video games have been popular for decades (Egenfeldt-Nielsen et al., 2013; Gee, 2003; Ivory, 2015; Kent, 2010; Wolf, 2015). Performance in first-person shooter (FPS) video games relies on acquired perceptual and motor skills (Green and Bavelier, 2003) and evidence suggests that playing these games enhances visuomotor and cognitive skills (Bavelier et al., 2012; Green and Bavelier, 2008) in a variety of visual and cognitive tasks (Colzato et al., 2013; Dye et al., 2009; Green and Bavelier, 2003, 2007) in children and adolescents (Adachi and Willoughby, 2013b,c,a; Funk and Buchman, 1996) as well as adults (Green and Bavelier, 2006, 2007; Kowal et al., 2018). There is, consequently, much interest in research on this topic, including the potential applications in digital therapeutics (Hong et al., 2021), but there is a paucity of studies on gaming performance itself.

Despite the rapidly growing popularity of competitive esports and the years spent by competitive and professional players to advance their skills (Popper, 2013), the field continues to lack established and objective benchmarks for individual skill. Many of the commonly used performance metrics in esports are unreliable measures of individual skill (Pedraza-Ramirez et al., 2020). Particularly in team games, key benchmarks (e.g., kill-death ratio, damage dealt, win-rate) confound the ability of an individual player with that of either their teammates, the opposing team, or the coordination of the team as a whole (Voida et al., 2010). For instance, a player with mediocre skill is still capable of obtaining a high rank simply by playing with individuals of a much higher skill level. The opposite is true for a great player with lesser-skilled teammates. Reliable, repeatable, and objective individual skill assessments with both intra- and inter-individual comparisons, on both short and longitudinal timescales, are needed to characterize true player rankings, efficacy of training, and to evaluate the impact of hardware, software, and player health (e.g., posture, exercise, sleep, diet, and dietary supplements) on performance.

Visuomotor psychophysics offers experimental approaches that are well-suited to assess individual players’ FPS performance. Successful FPS play requires efficient identification and localization of relevant visual stimuli, and dynamic movements followed by well-timed shot responses (Pluss et al., 2020). One fundamental facet of FPS play is “flicking” – the action of rapidly moving the cross-hair to an object or enemy and firing a shot to damage or destroy the target. When playing with a computer mouse and keyboard, which is typical for expert players, flicking can be achieved by reach movements when targets are far away, or wrist and finger movements when targets are nearby. Protocols for measuring such ballistic kinematics are well established, and have been applied in a broad range of cognitive and visuomotor tasks (Desmurget and Grafton, 2000). As of yet, specific knowledge regarding FPS performance, based on rigorous visuomotor psychophysics models, remains scarce. Visuomotor skills are specialized, and are often constrained to the contexts and modalities in which they are learned (Fahle, 2005; Heuer and Hegele, 2008; Maniglia and Seitz, 2018). Thus, current laboratory-based visuomotor psychophysics research provides limited knowledge about FPS performance.

Motor acuity is defined as the ability to execute actions more precisely, and within a shorter amount of time (McDougle and Taylor, 2019; Mü ller and Sternad, 2004; Shmuelof et al., 2012, 2014; Wilterson, 2021). There is a paucity of studies examining motor acuity, a gap likely linked to the tight resource constraints on laboratory-based studies. The handful of lab studies that examine motor acuity have used relatively simple motor tasks (Flatters et al., 2014), like drawing circles as fast as possible within a predefined boundary (Shmuelof et al., 2012), throwing darts (Martin et al., 1996), or center-out reaching and grasping (Jordan and Rumelhart, 1992; Shadmehr and Mussa-Ivaldi, 1994). Our group previously used a large sample of Aim Lab™ performance data (over 7,000 players and over 60,000 repeats of the 60 second Gridshot task) over a period of months to examine motor learning (Listman et al., 2021), using hits per second as a proxy for motor acuity. Here, we propose a new approach for measuring motor acuity and calculating flicking skill by characterizing an individual player’s speed-accuracy tradeoff, which we call the Flicking Skill Assessment (FSA).

It is both well established in the literature and widely discussed in the competitive gaming community that human performance exhibits a speed-accuracy tradeoff (SAT) (Heitz, 2014; Pluss et al., 2020): the speed of a response or action is negatively correlated with the accuracy or precision of that action. Players can be very fast and less accurate, very accurate and slow, or somewhere in between. This effect can be evident for different aspects of speed (e.g., reaction time, movement speed) and different aspects of accuracy (e.g., percent of correct responses/decisions, movement accuracy and variability). A hallmark of SAT is the ability to adapt to current demands and prioritize speed and accuracy relative to each other. If a task requires very fast responses or movements, a player may sacrifice accuracy to maximize speed. If there is a high cost to incorrect responses, a person may take longer to respond or move more slowly to maximize accuracy. In FPS performance, one way to characterize SAT is in terms of shot behavior, with the spatial error of shots representing accuracy and the number of shots fired per second representing speed. Performance on a single task with a single incentive for speed vs. accuracy is insufficient to estimate a player’s flicking ability, as two players with the same ability may choose a different strategy in managing the SAT. Thus, ability cannot be characterized from a single data point, since ability and strategy would be confounded. The FSA assesses performance in a plurality of conditions in which priorities or incentives for speed and accuracy differ, to isolate an individual player’s skill from their chosen trade-off between speed and accuracy. The result of this assessment is a measure of motor acuity for quantifying skill in FPS flicking tasks, which is independent of bias or strategy.

We used Aim Lab™, an FPS video game that assesses and trains players to optimize their performance, i.e., an automated personal trainer. Aim Lab™ comprises a wide range of tasks that replicate gaming scenarios, each matched with a particular data analysis procedure and inspired by visuomotor psychophysics. We recruited professional esports athletes (specializing in several different game titles and roles within a team) to play two tasks, one of which incentivized speed and the other of which incentivized precision. In addition to assessing SAT and motor acuity via shot behavior, we also characterized kinematics of players’ mouse movements. Players often aim for and destroy a target by initiating a movement towards the target, increasing and subsequently decreasing movement speed, then firing a shot after slowing down or coming to a full stop. We characterized these movements by fitting a sigmoid to the time-series of mouse positions, with the best-fit parameter values indicating a movement’s kinematics (i.e., reaction time to initiate a movement, movement speed, movement accuracy, variability or precision of a plurality of movements). In addition to these typical flick-and-land movements, players may choose to maximize speed by shooting “on the fly” instead of slowing down before firing. This type of movement behavior is referred to as a “swipe” movement, and we developed an approach to characterize the extent to which movements resembled a swipe versus a typical flick-and-land movement. We found that movement kinematics were shaped by task demands. In particular, reaction times were shorter and movements were less precise and more “swipey” for the task that incentivized speed over precision. Furthermore, we found that individual differences in movement efficiency (number of movements needed to hit a target) and movement kinematics (reaction time, precision, and swipiness) were highly predictive of motor acuity; thus, revealing the aspects of movement behavior that contribute to a player’s skill rank.

The FSA, our method for measuring motor acuity, may be used to assess and compare skill between and within players, as well as for training to improve skill and tracking improvements over time. The flexibility, accessibility, and engaging nature of FPS games also makes the FSA a promising tool for other applications, such as motor rehabilitation and monitoring and training cognitive health and fitness.

## 2 METHODS

### 2.1 Participants

Performance data from 32 professional and semi-professional male esports players (mean age = 22.47 *±* 3.62) were collected, specializing in different game titles: 4 Valorant players (Riot Games, 2020), 10 PUBG: Battlegrounds players (PUBG Studios, 2017), and 18 Rainbow Six Siege players (Ubisoft, 2015). Data were acquired initially for commercial purposes, stored separately from player account data and without personal identifiers. For this study, the data were reanalyzed post-hoc and informed consent was not required (Advarra Institutional Review Board).

### 2.2 Apparatus

Aim Lab™ is a commercial software product written in the C# programming language using Unity game engine (Helgason et al., 2005). Unity is a cross-platform video game engine used for developing digital games for computers, mobile devices, and gaming consoles (Brookes et al., 2020). Players download Aim Lab™ directly to their desktop or laptop PC. Players control their virtual weapon in Aim Lab™ tasks using a mouse and keyboard, while viewing the game on a computer screen.

The participants each completed the tasks remotely using their own gaming set-up. This included their own hardware such as the PC, monitor, mouse and mouse pad. In addition, there were other individual settings such as display size, viewing distance, chair height, and mouse counts per inch (CPI). We assumed that there was a wide range of equipment combinations amongst the participants, and we did not control for any differences in gaming set-ups between players; however, mouse acceleration was disabled. The orientation of the player’s view of the environment (controlled by the player’s mouse) was recorded in Euler angles and sampled at 120 Hz, then were uploaded to our secure servers. The crosshair, marked with a dot, was placed always at the center of the screen and corresponded to the direction in which a shot would be fired. When the player clicked to left mouse button to shoot, the projectile would move in a straight line pointing away from the player’s virtual avatar.

### 2.3 Task Descriptions

Aim Lab™ includes a variety of different task scenarios for skill assessment and training, each tailored to a facet of FPS play. These task scenarios assess and train a number of psychophysical processes, including: visual detection, motor control, tracking moving targets, auditory spatial-localization, change detection, working memory capacity, cognitive control, divided attention, and decision making. Each task can be customized to prioritize accuracy, speed, or any basic component of performance over others. During every round, players are granted points for each target that they successfully track or shoot and destroy. Additional points are rewarded for targets destroyed more quickly or tracked for a longer period. Players attempt to maximize their score on each round by destroying or tracking as many targets as possible.

In this study, we used task scenarios that assessed the players’ flicking skill, a combination of visual detection, motor planning, and motor execution. Specifically, we used two tasks, Gridshot and Sixshot, that were very similar but differed in target size to characterize each player’s SAT and estimate their motor acuity. The tasks were designed to incentivize players to maximize their score by either prioritizing accuracy over speed when targets were smaller, or prioritizing speed over accuracy when targets were larger. Each of the 32 professional-level esports athletes played 6 runs of each of the two tasks.

#### 2.3.1 Gridshot

In Gridshot (Fig. 1A) there are 3 targets presented simultaneously at any given time, with a new target appearing (spawning) once an existing target is destroyed. All targets are the same size, ranging between 1.3°and 1.7°(degree of visual angle), assuming a range of viewing distances and a range of values for the field of view in the virtual environment of the game (set by the player). Spawn locations are randomized to 1 of 25 positions in a 5 *×* 5 grid, ranging between 4.8°and 9.1°wide and 5.1°and 7.8°high and similarly depending on viewing distance and field of view. The player destroys a target by moving their mouse to aim then clicking the left mouse button to shoot. Due to multiple targets being present at once, and combined with the unlimited target duration and no explicit incentive to destroy any specific target, the players themselves must decide the order in which to destroy the targets. Players receive immediate feedback upon target destruction; the game emits an explosion sound and the orb-shaped target splinters into multiple pieces which then disappear. Players receive a summary of performance feedback after each 60 second run of Gridshot. These summary metrics include score, hits per second (number of targets successfully destroyed per second), and hit rate (percentage of shot attempts that successfully hit a target). Points are added to the score when the targets are hit and, in contrast, points are subtracted for shots that missed the target. The player’s score is displayed at the top of the screen throughout the run, automatically calculated in the game’s software, and sent to a secure server. The number of points added for each target destroyed is scaled by the time since the previously destroyed target. In other words, the shorter the time it takes to destroy the next target relative to when the previous target was destroyed, the more points are added to the player’s score. Thus, players are incentivized to quickly plan their next movement and shoot targets rapidly. Even though players are shown multiple metrics at the end of each run, it is likely that they are consciously optimizing for increased score during the actual runtime of the task.

**Figure 1.**
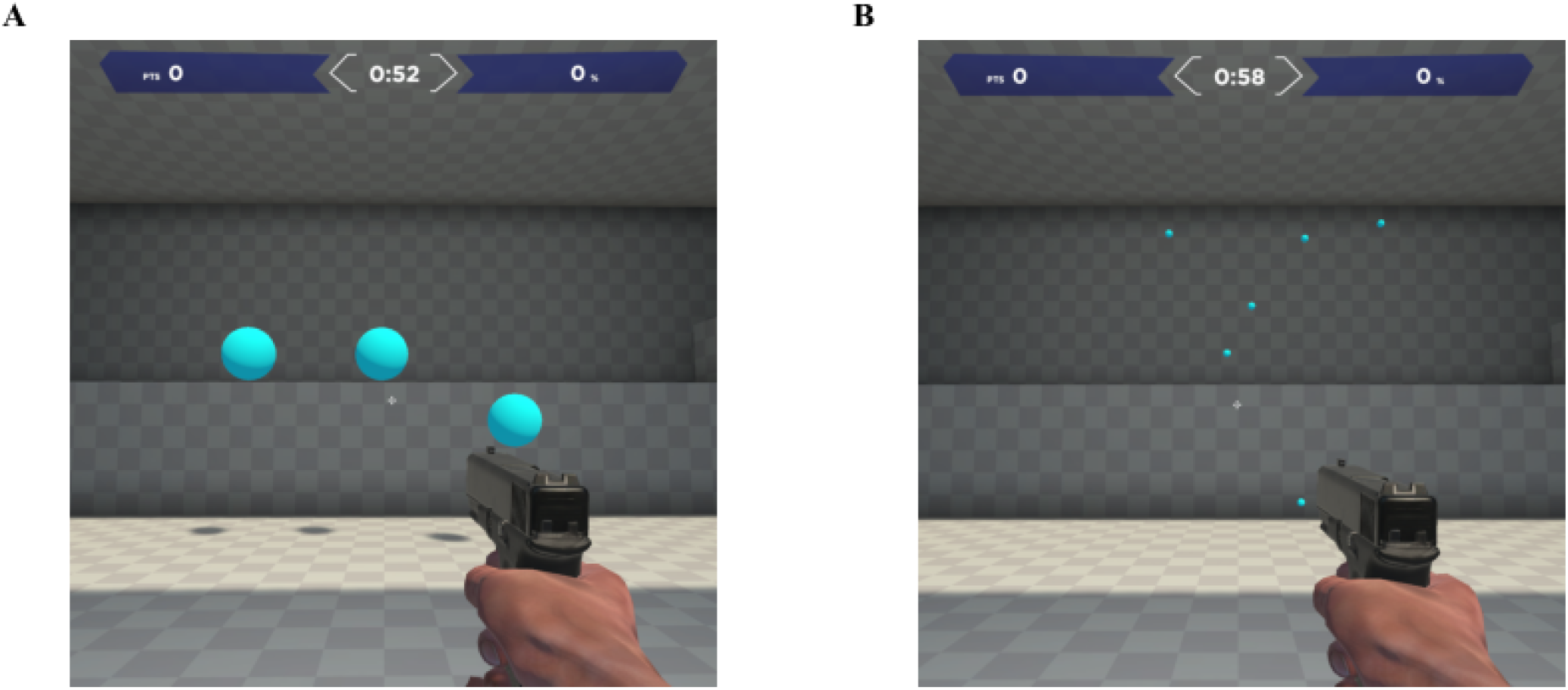
Screenshots from Aim Lab™, displaying a single frame from the: **A)** Gridshot, and **B)** Sixshot tasks.

#### 2.3.2 Sixshot

Sixshot (Fig. 1B) is very similar to Gridshot, whereby they share identical possible target locations. However, the Sixshot task bears the following differences to Gridshot: 6 targets are present at a time (rather than 3), and the targets are approximately 14% the size of Gridshot targets. If a player is maximizing for score, the smaller target size demands much greater shot accuracy from the player than the targets in Gridshot.

### 2.4 Data Analysis

#### 2.4.1 Mouse calibration

To measure the player’s physical mouse movements, we converted the player’s orientation in the virtual environment to the corresponding movements of the mouse on the mouse pad, which has units of centimetres (cm). This required additional information about the relationship between physical mouse movement and changes in player orientation, which varied across players due to their settings in Aim Lab™, and their own hardware and software. For each player, we recorded their in-game settings that governed the field of view and their mouse sensitivity – the magnitude of a change in camera rotation (in degrees, °) derived from a single increment or count of the mouse hardware. The individual mouse sensitivities were constant across x and y for each player. Each player’s mouse (either hardware, software, or both) had a unique setting that determined the number of counts that resulted for one inch of distance traveled (counts per inch, CPI).

Player camera orientation was converted to physical mouse movements as detailed below:

1. Mouse sensitivity × 0.05 = angle increment (degrees turned per count)
2. Total degrees turned / angle increment = counts
3. (Counts / CPI) × 2.54 = physical distance traveled (cm)

The value of 0.05 is an arbitrary constant used in the Unity software code to scale mouse counts to degree increments. After this conversion, across all players and tasks the maximum target distance was 5.60 cm. Thus, all targets were proximal enough that players could have landed on every target using a combination of only wrist rotations and bending or straightening the fingers, i.e. forearm or elbow movement were not strictly necessary.

#### 2.4.2 Movement parsing

To assess movement kinematics, we first parsed the time-series of each player’s orientation in the virtual environment of the game, as controlled by their mouse movements. Each time-point (or sample) was labeled as either in-motion or stationary. Epochs were labelled as in-motion after several consecutive samples exceeded a velocity threshold. Conversely, epochs were labelled as stationary after several consecutive samples fell below a velocity threshold. The number of consecutive samples used and the values of the thresholds were determined using a proprietary algorithm, procured from a large dataset of Aim Lab™ players.

The first movement a player makes towards a target is known as a primary movement. During each sixty-second run of the tasks, these primary movements were often followed by a corrective movement, i.e., an action in response to not having destroyed the target with the primary movement. For instance, the primary movement could be hypermetric, indicating that the player’s crosshair passes beyond the target and they must initiate a corrective movement in the opposite direction back towards the target. Alternatively, the primary movement could instead be hypometric, falling short of the target and requiring a corrective movement in the same direction as the primary movement. At times, multiple corrective movements were needed to destroy the target. Each movement component was fit separately, whether it was a primary or corrective movement.

#### 2.4.3 Movement kinematics

By fitting parametric functions to the mouse movement data, we quantified the features of an individual’s motor behavior. This resulted in measurements of speed, precision, accuracy, reaction time (henceforth shortened to SPAR), as well as swipiness (see Section 2.4.4). Specifically, player orientation during each period labeled as in-motion was fit with a sigmoidal function:

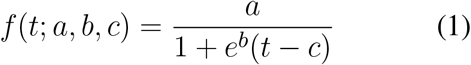

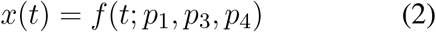

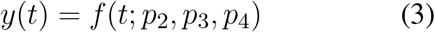

where Eq. 1 defines the sigmoid. The values of *x(t)* in Eq. 2 represent a model of the horizontal component (rotation about the y-axis) of the movement trajectory for each time-point sample. The values of *y(t)* in Eq. 3 represent a model of the vertical component (rotation about the x-axis) of the movement trajectory for each time-point sample. Example illustrations of the model components *x(t)* and *y(t)* are shown in Fig. 2. The top row of Fig. 2 shows two example movements in units of centimeters, transformed using the mouse calibration. The bottom row of Fig. 2 shows the same two example movements, scaled to normalized units so that 1 corresponds to the target location. The left column of Fig. 2 shows an example of a flick-and-land. In these panels, the shot (vertical dashed line) occurs after the movement ends and the movement lands at the target location (horizontal dotted line). The right column of Fig. 2 shows an example of a swipe. The shot (vertical dashed line) is made during the middle of the movement, and the movement lands well past the target location (horizontal dotted line). The values of the parameters (*p*_*1*_, *p*_*2*_, *p*_*3*_, *p*_*4*_) were fit to each individual movement trajectory using the Levenberg-Marquardt algorithm (Levenberg, 1944).

**Figure 2.**
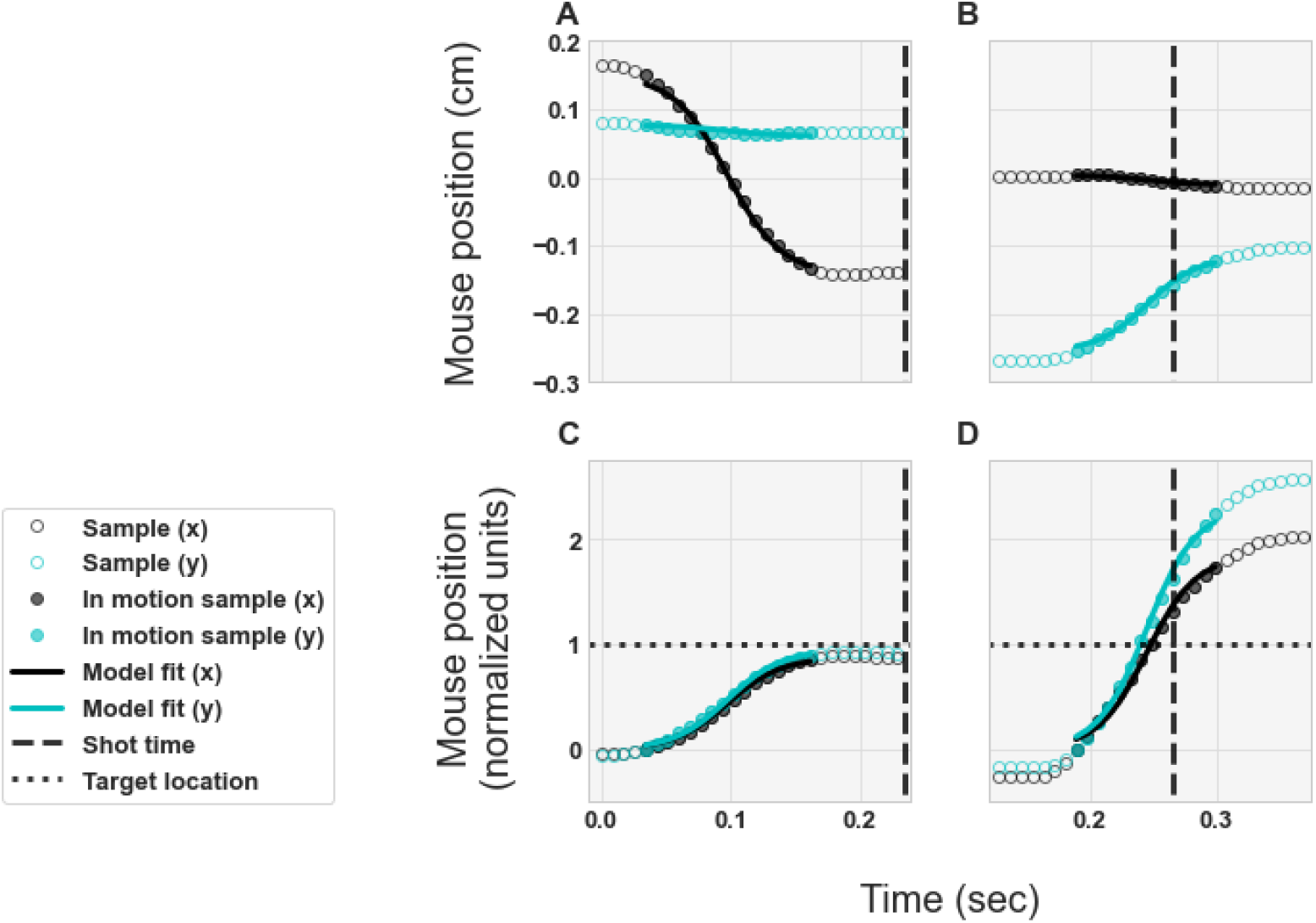
Examples of the movement trajectories and model fits. **A, C)** Example flick-and-land. **B, D)** Example swipe. **A, B)** Physical movements in units of centimeters. **C, D)** normalized units -so that 1 corresponds to target location. Circles illustrate the samples of the player’s movement trajectory. Curves illustrate the models of the movement trajectories, as expressed by Eqs. 1, 2, and 3, with best-fit values for the parameters.

In Eqs. 2 and 3, the x- and y-movement components were fit with shared parameters *p*_*3*_ and *p*_*4*_.

SPAR metrics were then calculated from the best-fit parameter values:

1. Speed (cm / second): peak speed at the midpoint of the movement.
2. Precision (1 / median absolute % distance to the target): variability in the accuracy across trials/targets.
3. Accuracy (% distance to the target): distance between the landing location of the movement and the center of the target.
4. Reaction time (sec): time interval between when the target appeared and the initiation of the movement, i.e., when the movement reached 5% of its endpoint.

Specifically, the movement speed and accuracy of each parsed movement were quantified as:

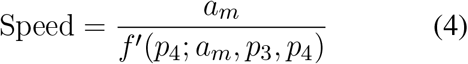

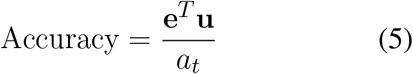

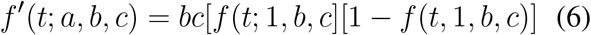

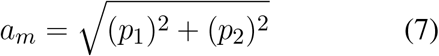

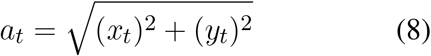

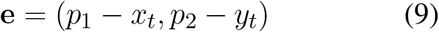

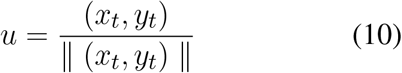

The function *f’(t)* is the derivative of the sigmoidal function (Eq. 6). The value of *a*_*m*_ represents the amplitude of the movement (Eq. 7) and the value of *a*_*t*_ represents the distance to the target location (Eq. 8). The vector **e** represents the movement error (Eq. 9) and the vector *u* represents a unit vector in the direction of the target location (Eq. 10). Movement speed was re-scaled using the mouse calibration protocol (to have units of: cm / second, see Section 2.4.1). Precision was calculated as the variability in accuracy, using a robust measure of variability (the median of the absolute difference from the target center, rather than the standard deviation) to minimize the impact of outliers.

#### 2.4.4 Swipiness

To characterize the degree to which each ballistic movement resembled a swipe (the action of shooting on the fly) versus a flick-and-land (slowing down and stopping before firing) movement, we compared the time of each shot that a player made with the time of the midpoint from the associated ballistic movement. An ideal swipe corresponds to firing a shot at the midpoint of a movement, i.e., the time point with maximum speed. On the other hand, an ideal flick-and-land corresponds firing a shot only after the movement has ended, long after the midpoint of the movement. Thus, swipiness was calculated from dividing the time of the shot by the time of the midpoint of the movement (*p*_*4*_), divided by 2. This resulted in a swipiness value of 0.5 (arbitrary units) for an ideal swipe, and a swipiness value *≥* 1 for an ideal flick-and-land. There was no swipiness value computed for a movement that had no associated shot, i.e., if there was no shot between the initiation of one movement and the initiation of the next movement.

Swipiness itself is a granular measure of shot speed as it relates to the movement trajectory, with lower swipiness indicating that the firing of a shot occured earlier in the trajectory. Additionally, the absence of a swipiness value indicates that a movement was not associated with a shot. Across trials within a certain context or task, the number of movements with no swipiness value reflects the need for players to make multiple movements to destroy targets.

#### 2.4.5 Data post-processing

For each player, we computed SPAR for every movement in Gridshot and Sixshot and swipiness for every movement that had a corresponding shot fired. We then applied several steps of post-processing.

Firstly, an upper and lower band threshold was applied to remove outliers (presumed to be failures in the trial parsing or SPAR fits). For all of the metrics, the 95th percentile was used for the upper band value and 0 was used for the lower band value. Sigmoid fits with an r-squared value of less than 0.5 were pruned. Each accuracy value was multiplied by 100, to convert from proportion to percent of the distance to the target. We then subtracted 100 from each value such that hypometric movements had negative values and hypermetric had positive values, i.e., accuracy values of 0 indicate that the player fired a shot directly at the centre of the target to destroy it. We computed the median of each metric, separately for each individual player, and separately for each task and movement-type condition. This yielded a total of 20 variables per player: 5 metrics (median speed, precision, accuracy, reaction time, and swipiness) × 2 movement types (primary versus corrective) × 2 tasks (Gridshot versus Sixshot).

A z-score was calculated for each metric across players, combining both Gridshot and Sixshot (for each metric and movement type) to permit statistical comparisons between the tasks. The distributions of these data are shown in Fig. 3. Once z-scored, data was tested for normality using Kolmogorov-Smirnov; the resulting *p*-values indicated that the majority of metric distributions were non-normal (6 out of 10 variables where *p*s *<* 0.05) and therefore non-parametric approaches to statistical analysis were taken. To mitigate the confounding effects of mouse sensitivity, this vector was converted into a diagonal matrix and regressed out of both the movement kinematics and motor acuity. To do so, we expressed the z-scored movement kinematic metrics **Y** (a 32 × 20 matrix: 32 players and 20 kinematic variables, as stated above) as a linear prediction of the z-scored mouse sensitivities **A** (a 32 × 32 diagonal matrix) multiplied by regression coefficients **X**:

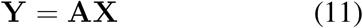

**Figure 3.**
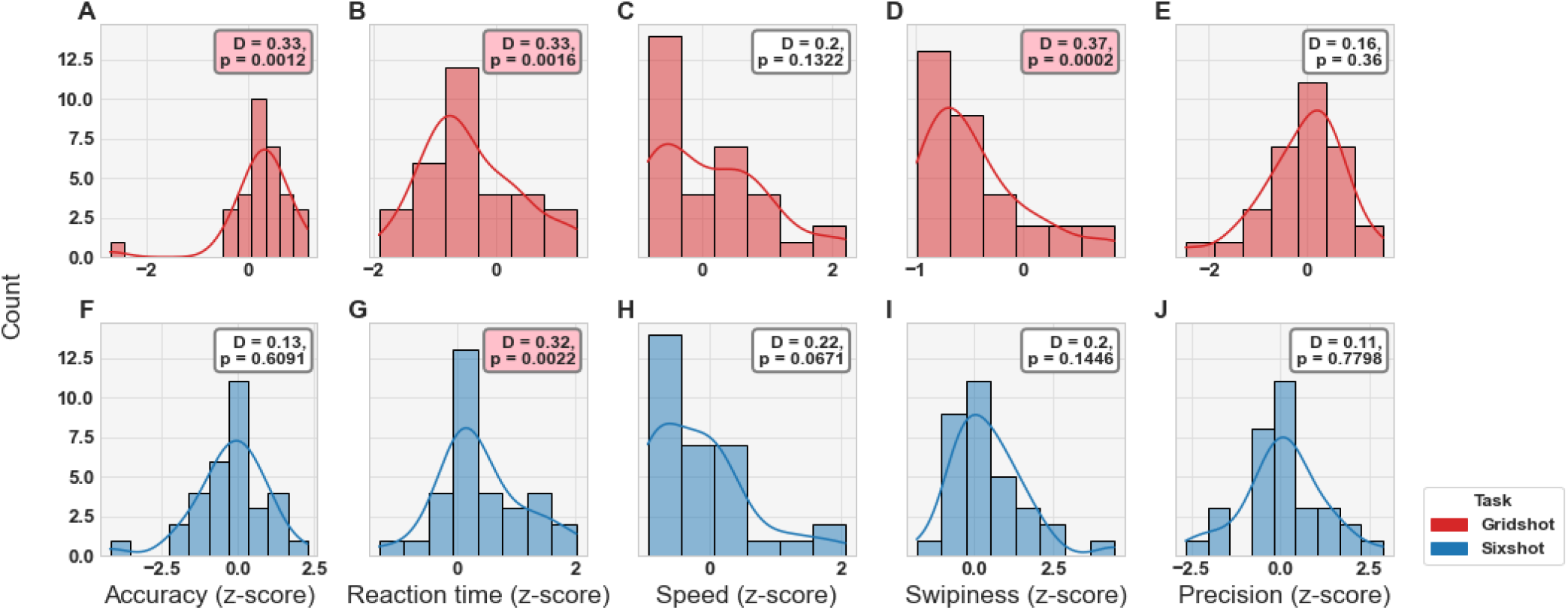
Distribution plots for each metric by task (see Legend). **A–E)** Gridshot. **F–J)** Sixshot. Statistical results from Kolmogorov-Smirnov test for normality displayed in annotation boxes; where D = KS-statistic and p = *p*-value. Pink annotation boxes indicate a *p*-value *<* 0.05.

First, the inverse of the mouse sensitivity diagonal matrix **A**^**#**^ was calculated and multiplied by the movement kinematics matrix **Y** to estimate the regression coefficients 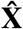:

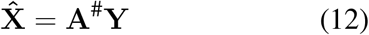

Second, we computed the components of the movement kinematics that could be predicted by the mouse sensitivities **Ŷ** by multiplying 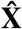 with the mouse sensitivity diagonal matrix **A**:

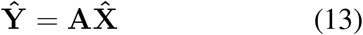

Third, the residual movement kinematics matrix, **R**, was computed by subtracting **Ŷ** from **Y**:

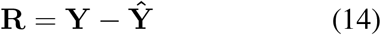

Mouse sensitivity was correlated with motor acuity and with primary movement reaction time (both tasks), speed (both tasks), and swipiness (Sixshot only). However, this correlation was removed by the regression procedure (Figs. 9-13, Supplemental Data).

## 3 RESULTS

### 3.1 Motor acuity

We predicted that players’ would exhibit a SAT as a function of the target size. In particular, we predicted that players would exhibit systematic differences in movement behavior between Gridshot and Sixshot. Gridshot’s large target size incentivizes players to maximize score by moving as quickly as possible with relatively low shot precision (1 / median shot error, as measured by the distance to the center of the nearest target in cm). In comparison, Sixshot targets are much smaller and require high precision, which consequently means that players are motivated to slow down in order to maximum their score.

Fig. 4 illustrates the SAT and the Flicking Skill Assessment (FSA). The left panel of Fig. 4 shows shot speed as a function of the median shot error for Gridshot and Sixshot. Each line connects the Gridshot and Sixshot data for a single player, portrayed respectively by diamonds and squares. Our findings indicate that each player followed the expected pattern in behavior: lower precision and more total shots in Gridshot compared to Sixshot. This demonstrates that players responded strategically and appropriately to the different demands of the two tasks. The right panel of Fig. 4 re-plots these results in terms of shot precision (1 / shot error). The curves represent the transformed lines from the left panel.

**Figure 4.**
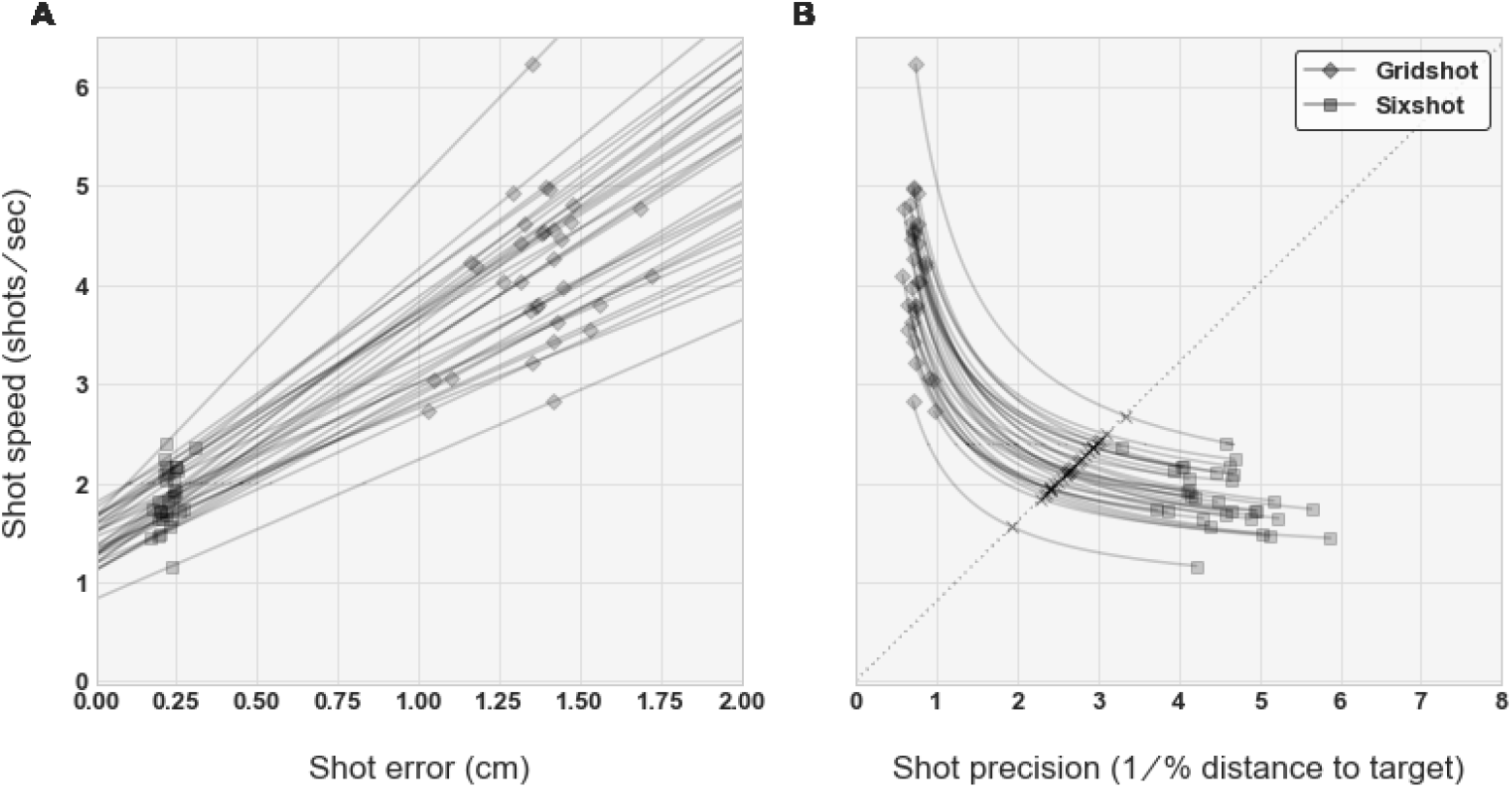
**A)** Shot speed (shots per second) versus shot error. Solid lines illustrate the line of best fit for each player by task (see Legend). **B)** SAT curves. Dashed line spans from (0, 0) to three-times the standard deviation, plus the mean of the distribution of shot precision (x-axis) and shots per second (y-axis). The intersection of individual curves with the diagonal, marked with “×”, indicates motor acuity (arbitrary units).

The dashed diagonal line spans from (0, 0) to three-times the standard deviation, plus the mean of the shot precision distribution (x-axis) and shots per second distribution (y-axis). This line represents the axis of motor acuity; SAT curves placed towards the upper right indicate better performance, i.e. higher precision and faster speed. We obtained a flicking skill value for each player, by identifying the point along the diagonal line that intersects with that player’s SAT curve. Notably, these flicking skill values are related to certain aspects of movement kinematics, as detailed in the following subsections.

### 3.2 Movement frequency

Human flicking performance typically exhibits variability and error. At times, a primary movement will have very little error, while others will require subsequent corrective movements to successfully land on and shoot a target. For large targets, such as those in Gridshot, it is expected that the proportion of targets destroyed which required a corrective movement would be lower. This is because even with large movement error, primary movements are more likely to land on the target. Contrarily, players will often have to make corrective movements to accommodate for the targets being much smaller, such as those in Sixshot.

For both tasks, Fig. 5 shows the percentage of targets that were destroyed with a single primary movement or with a double movement of both primary and corrective. With the exception of 3 players, the results reflect the pattern we predicted. The number of single movements to destroy the target was greater in Gridshot, and double movements to destroy the target were more frequent in Sixshot. The number of movements per target destroyed was related to player skill, namely with motor acuity (after regressing out mouse sensitivity). Fig. 6 shows motor acuity on the x-axis and proportion of targets destroyed following a single movement on the y-axis for Gridshot (Fig. 6A), and Sixshot (Fig. 6B). Flicking skill was positively correlated with the proportion of targets that were destroyed with a single movement (*p* = 4.19e-04) in Gridshot. However, there was no significant correlation in either direction for Sixshot (*p* = 0.26). This suggests that players with a higher motor acuity were more likely to require a single shot to destroy a target in Gridshot, and therefore were more efficient with their movements.

**Figure 5.**
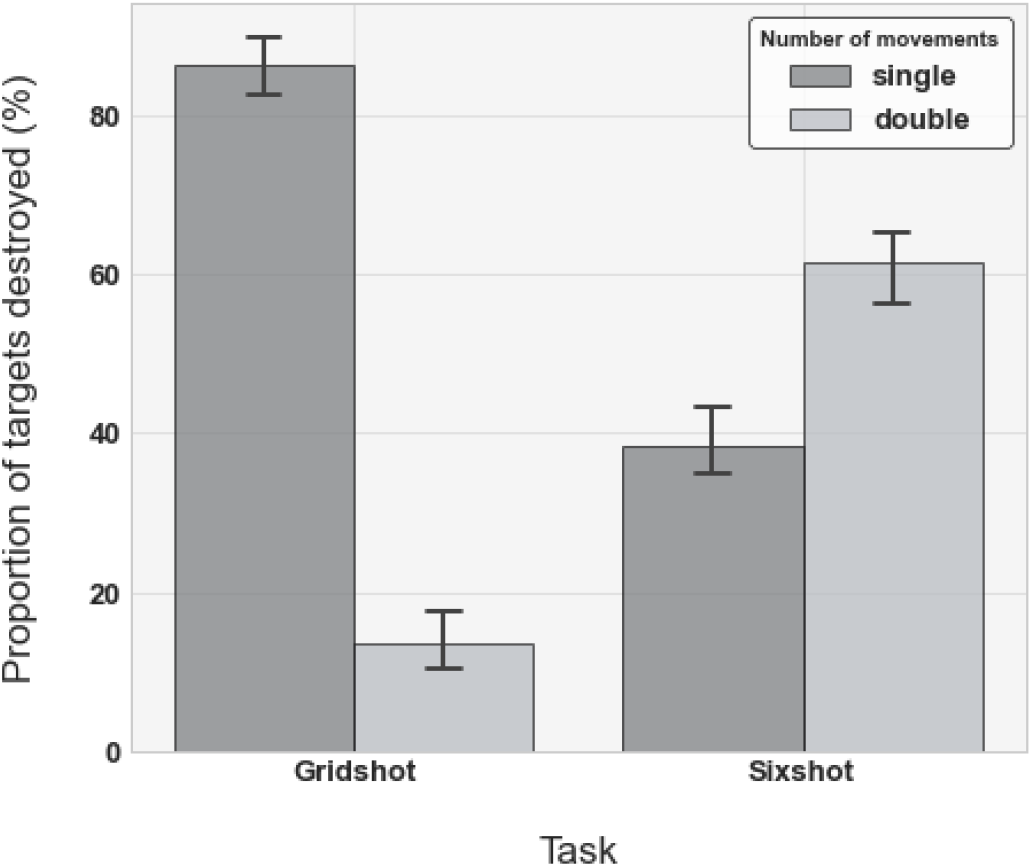
The mean proportion of single movements associated with each target per task (see Legend). Error bars illustrate the standard deviation across all players.

**Figure 6.**
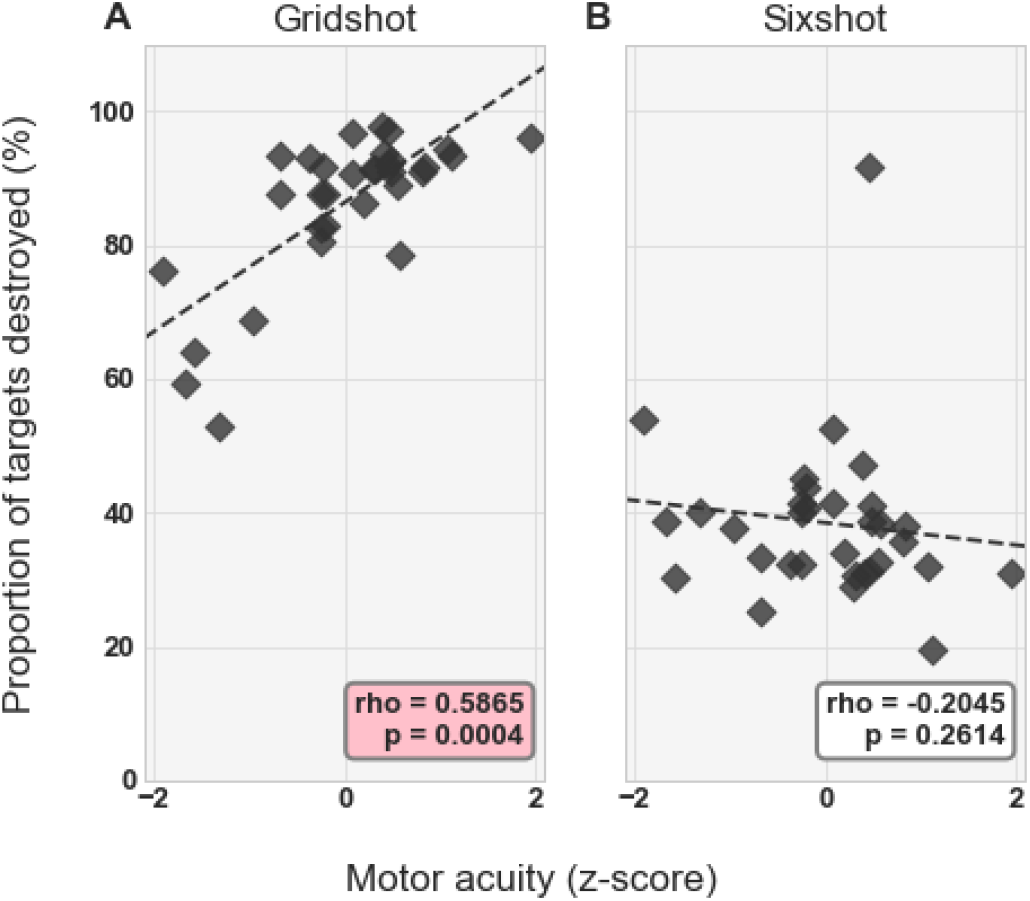
Correlation between the proportion of single movements and motor acuity: **A)** Gridshot. **B)** Sixshot. Statistical results from Spearman’s Rank Correlation displayed in annotation boxes; where rho = Spearman’s Correlation Coefficient and p = *p*-value. Pink annotation boxes indicate a *p*-value *<* 0.05.

### 3.3 Task-dependence of movement kinematics

Movement kinematics differed between the two tasks; Fig. 7 illustrates each player’s pattern of strategy change for each movement kinematic metric (after regressing out mouse sensitivity), with Gridshot on the x-axis and Sixshot on the y-axis. The results indicate that players were more hypometric, i.e., less accurate, in Sixshot compared to Gridshot (Accuracy; *p* = 8.85e-03). Furthermore, players were faster to react (Reaction time; *p* = 2.01e-05), move their mouse at a faster pace (Speed; *p* = 1.00e-06), and fire earlier in their movement trajectory (Swipiness; *p* = 5.81e-05) in Gridshot compared to Sixshot. There was no statistical evidence of a significant difference in the variability of accuracy between tasks (Precision; *p* = 0.29), indicating that players were similarly consistent in their landing positions for Gridshot and Sixshot.

**Figure 7.**
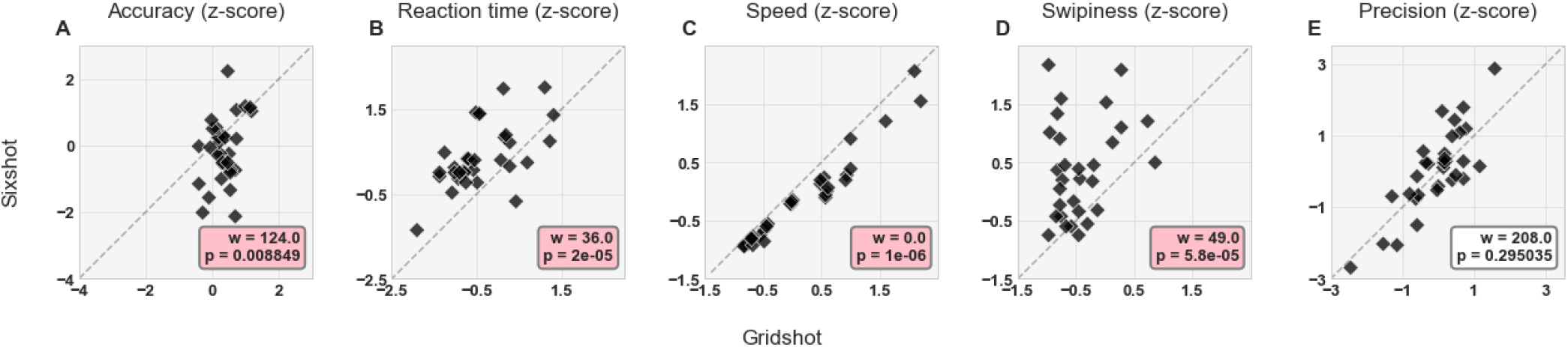
Differences in movement kinematics for Sixshot versus Gridshot: **A)** Accuracy. **B)** Reaction time. **C)** Speed. **D)** Swipiness. **E)** Precision. Each data point within each panel corresponds to a different player. Statistical results from the paired Wilcoxon signed-rank test displayed in annotation boxes; where w = sum of ranks, and p = *p*-value. Pink annotation boxes indicate a *p*-value *<* 0.05. The dashed diagonal represents the values at which the kinematic z-scores for Gridshot is equal to Sixshot.

In this analysis, we focused on the metrics for primary movements only; those of which cover the most distance to the target. Nevertheless, our approach in measuring movement kinematics can be used for both primary movements and corrective movements. The results above (Fig. 5) suggest that the players’ objectives were to destroy targets with as few movements as possible. Primary movements encompass the quality of their motor ability to do so, whereas corrective movements are strongly influenced by the movement(s) that preceded them towards the same target. Therefore, primary movements provide a more straight-forward interpretation to the kinematics compared to their corrective counterpart.

### 3.4 Individual differences in motor acuity and kinematics

Motor acuity, as measured with the FSA, was predictive of individual differences in movement kinematics. Fig. 8 shows the relationship between individual differences in motor acuity and movement kinematics (after regressing out mouse sensitivity). For Gridshot, motor acuity was negatively correlated with reaction time (*p* = 5.45e-03) and swipiness (*p* = 2.91e-02), while positively correlating with precision (*p* = 1.71e-04). For Sixshot, motor acuity was also negatively correlated with reaction time (*p* = 6.10e-04) and positively with precision (*p* = 1.30e-02). For both tasks, the interpretation of these results are that players with greater motor acuity initiated movements more quickly and landed with lower variability. Moreover, in Gridshot only, these players were also found to fire shots earlier in the movement trajectory.

**Figure 8.**
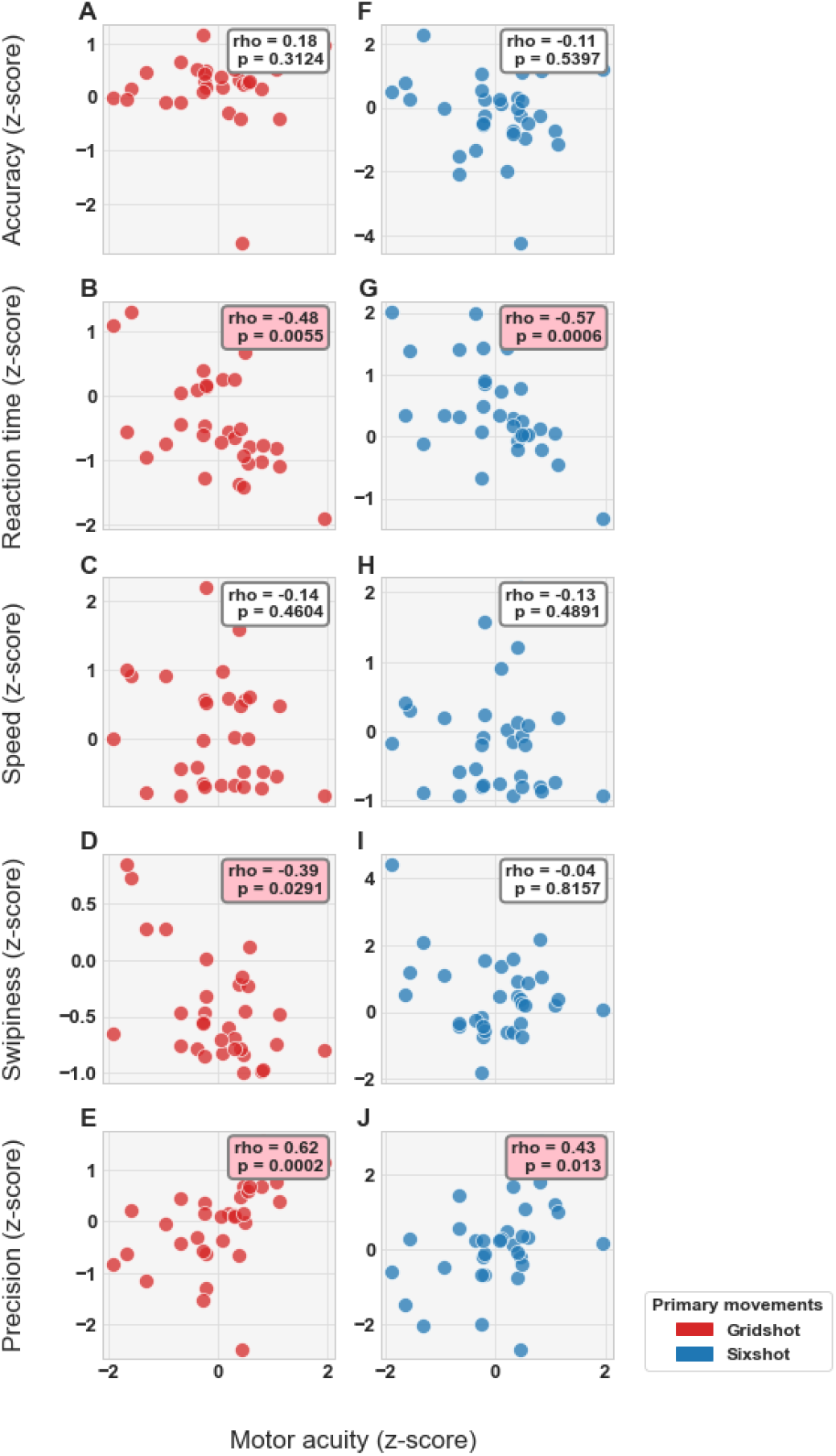
Correlation between player kinematics and motor acuity for each metric in each task (see Legend): **A–E)** Gridshot. **F–J)** Sixshot. Statistical results from Spearman’s Rank Correlation displayed in annotation boxes; where rho = Spearman’s Correlation Coefficient and p = *p*-value. Pink annotation boxes indicate a *p*-value *<* 0.05.

## 4 DISCUSSION & CONCLUSION

In this study, we developed and validated the flicking skill assessment (FSA), a novel approach to objectively assess individual player skill in FPS games. Furthermore, we elucidated the systematic relationship between motor acuity (measured by characterizing speed-accuracy tradeoff, SAT, curves) and movement kinematics. Our results reveal the individual differences in motor acuity and movement kinematics between professional-level FPS players, as well as context-dependent performance differences within individual players according to task demands. We also show that, for comparable data across players, it is necessary to account for differences in mouse sensitivity settings among players and to transform mouse and target location data points from orientation in the virtual environment to centimeters on the mouse pad. The proposed FSA is an elegant and efficient approach which requires only an adjustment of target size between similar tasks.

The players’ performance were found to differ between tasks, whereby players used a more conservative strategy in Sixshot compared to Gridshot. This is evident in the players’ shot behavior, with fewer shots and smaller spatial error when targets are very small (Sixshot) and a greater number of shots and greater spatial error when targets are large (Gridshot). Auxiliary to the players’ shot behavior, the movement kinematics provided detailed information about the nature of this shift in strategy between tasks: players take longer to plan their movements, move slower, fire later in the trajectory, and are more likely to land hypometrically in Sixshot compared to Gridshot. Given that the targets in Sixshot are small and require less error compared to large targets in Gridshot, this adaption in shot and movement behavior is a rational and efficient approach. Players sacrifice speed for accuracy and display more of a bias towards conservative (hypometric) landing positions during Sixshot compared to Gridshot.

Our analysis characterized individual differences in motor acuity and movement kinematics across professional-level esports athletes. Regardless of task demands in a single context, players differ in the degree to which they can be both fast and accurate. Players with a higher motor acuity value, i.e., their SAT curves are further upwards and rightwards, were both faster and more accurate overall than players with a lower motor acuity value, and are thus considered to have a higher skill rank. Critically, motor acuity significantly correlated with a subset of movement kinematics. This demonstrates that individual differences in flicking skill are predictive of individual differences in kinematics. Greater motor acuity accompanied faster reaction times in both Gridshot and Sixshot, in addition to having greater precision and a tendency to shoot earlier in the trajectory (swipiness) in Gridshot (Fig. 8).

One crucial design element of the FSA is the inclusion of more than one task, each with distinct incentives regarding speed and precision. The placement of an individual’s entire SAT curve is the key to quantifying motor acuity, rather than the placement of a single data point. The SAT curve itself represents the possible combinations of shot speed and shot precision given the player’s skill, with the player’s current strategy determining where along the curve their current speed and precision is drawn from. Two players with identical SAT curves could have different strategies within the same task, such that their performance would sample different values for speed and precision on the same curve. Including multiple tasks, each with distinct incentives for speed and precision, enables the characterization of SAT across possible incentives, and thus isolates motor acuity.

Indeed, we have now implemented a version of the FSA with 3 different target sizes (Fig. 14). In this task, called Adaptive Reflexshot, one target at a time appears in a randomized location, confined to an imaginary ellipse in front of the player’s virtual avatar. Players have a limited time to destroy each target before it “times-out” and disappears. To provide a visual cue of time remaining, each target gradually becomes more transparent and then disappears. The target presentation duration is titrated according to performance: the duration is decreased (it becomes transparent more quickly) on the next trial after a target is destroyed and the duration is increased (it becomes transparent more slowly) on the next trial after a target time-out, separately for each target size.

A second crucial design element is the measurement of both speed and precision. It is not be possible to use a single performance metric as a proxy for motor acuity. For instance, retrieving only shot speed from both tasks can be confounded by players simply shooting faster with no ramifications for being inaccurate – a behavior that does not reflect high skill.

Our measure of motor acuity and the movement kinematics were derived from different behavioral sources: shot behavior and mouse movements, respectively. By identifying a systematic relationship between the two sources, our findings demonstrate that individual differences in player skill correspond to individual differences in underlying movement kinematics. We propose that the FSA, a measure of motor acuity, represents a powerful tool for quantifying skill in FPS flicking tasks. One of the major benefits of the FSA is the capability to isolate player ability from context-dependent strategy regarding SAT. Indeed, a player can achieve their best possible skill ranking by matching shot behavior with task incentives. However, their underlying ability will determine the upper limit of their measured motor acuity, i.e., when they adopt the optimal strategy across tasks in the FSA. Faced with different target sizes, players who are experienced and skilled will flexibly adjust their behavior to maximize performance. Shot speed and precision reveal their flicking ability, and the relative skill of individual players can be objectively compared by using this motor acuity metric.

Gamers are highly invested in optimizing their performance, including factors distinct from improving FPS skill per se. To that end, it is of great value for this community to establish objective benchmarks for how any given influence on performance can be altered or shaped to maximize gaming outcomes. Using the FSA, the industry could readily assess within-participant differences in motor acuity to reveal the influence or relative efficacy of particular hardware (e.g., mouse, mouse pad, monitor rendering latency), software (e.g., mouse settings, axes inversion), and various other factors related to the health and physical attributes of players (e.g., posture, exercise, sleep, diet, and dietary supplements). As the scientific study of gaming performance is in its infancy, the FSA is an unprecedented and broadly applicable tool to efficiently establish foundational knowledge in the field.

The FSA not only allows for comparisons to the population, but additionally provides feedback on an individual level. That is, each individual user may measure their own baseline during an initial session of the FSA so that repeated gameplay can monitor changes relative to this baseline. This is particularly salient for esports professionals, given that currently available metrics are unsubstantiated and team performance often clouds the details of individual performance (Voida et al., 2010). Elucidating the comparable strengths and weaknesses of each member of any given esports team would not only provide a basis for coaching, but for team composition as well.

This flexible tool is also a promising training regimen for improving FPS skill. As esports professionals, the participants involved in this study were able to recognize and adapt to the incentives for SAT in both tasks as expected. Players of a lower skill or experience level would be more likely to fail at adjusting their shot behavior according to task demands, causing performance to suffer. The FSA can readily identify such faulty or insufficient adjustments in strategy. To be specific, if the slope of the line fitted to a player’s shot speed by shot error is either negative, near zero, or near vertical, this indicates that the player did not make the trade-off between accuracy and speed to maximize performance between scenarios. An assessment or training protocol based on the FSA would be able to automatically provide actionable feedback: explaining the difference in incentives between the tasks based on SAT, and, for example, instructing the player to slow down or speed up depending on target size. Ultimately, repeated participation in the FSA could facilitate the improvement of flicking through experience and practice in conjuction with explicit instruction on the optimal strategies a player should use during play.

The ability of the FSA to provide, over a short period of time with no ceiling effect, a measure of motor acuity independent of strategy and SAT makes it a promising tool over a wide range of both basic and applied research; for example, visuomotor psychophysics and rehabilitation. As noted above (see Introduction), there is a paucity of academic studies examining motor acuity. The FSA has the potential to fill this critical gap in the study of human motor behavior and visuomotor psychopysics. As with any other instance of motor skill learning that is characterized by reinforcement learning [1], physical rehabilitation aims to improve motor acuity. At the same time, rehabilitation must also assess performance to provide motivating feedback for the patient and clinically relevant data for the therapist to guide rehabilitation.

Neuroplasticity drives motor learning, which itself depends on movement repetition and intensity (Nudo and Milliken, 1996; Nudo et al., 1996a,b). Neuroplasticity is also facilitated by active task engagement and enjoyment (Burdea, 2003; Burke et al., 2009b,a; Maclean et al., 2000, 2002; Plautz et al., 2000; Putrino et al., 2015; Winstein et al., 2016). Furthermore, calibrating task difficulty to an individual’s ability level is critical for rehabilitation (Wolf et al., 2006), because competency is an intrinsic motivator (Przybylski et al., 2010). The FSA satisfies all of these criteria: a challenging and engaging task based on repetitive movement behavior. Thus, incorporating the FSA into gamified rehabilitation therapy would enable objective quantification of behavioral or motor performance (e.g., kinematics, dynamics), and could be rapidly and inexpensively deployed at scale and remotely provided to large populations.

The participants in the current study were all experienced, highly-skilled FPS players. Further investigation is required to establish the degree to which our findings generalize to the broader population of FPS players (which spans a variety of skill and ability level), the general population, or the clinical populations. Additionally, it will be crucial to establish how lower-skilled players gradually improve through training and experience, and the corresponding changes in movement kinematics (e.g., shot behavior, SAT, etc.) as well as how to optimize training.

## AUTHOR CONTRIBUTIONS

ID, MS, and KD carried out data analyses and assisted with study design and manuscript preparation. DH and JL assisted with study design and manuscript preparation. WM designed and implemented the study apparatus and tasks.

## ACKNOWLEDGMENTS

The authors would like to thank the Aim Lab™ users whose playing time and performance data have contributed to this work.

## SUPPLEMENTAL DATA

**Figure 9.**
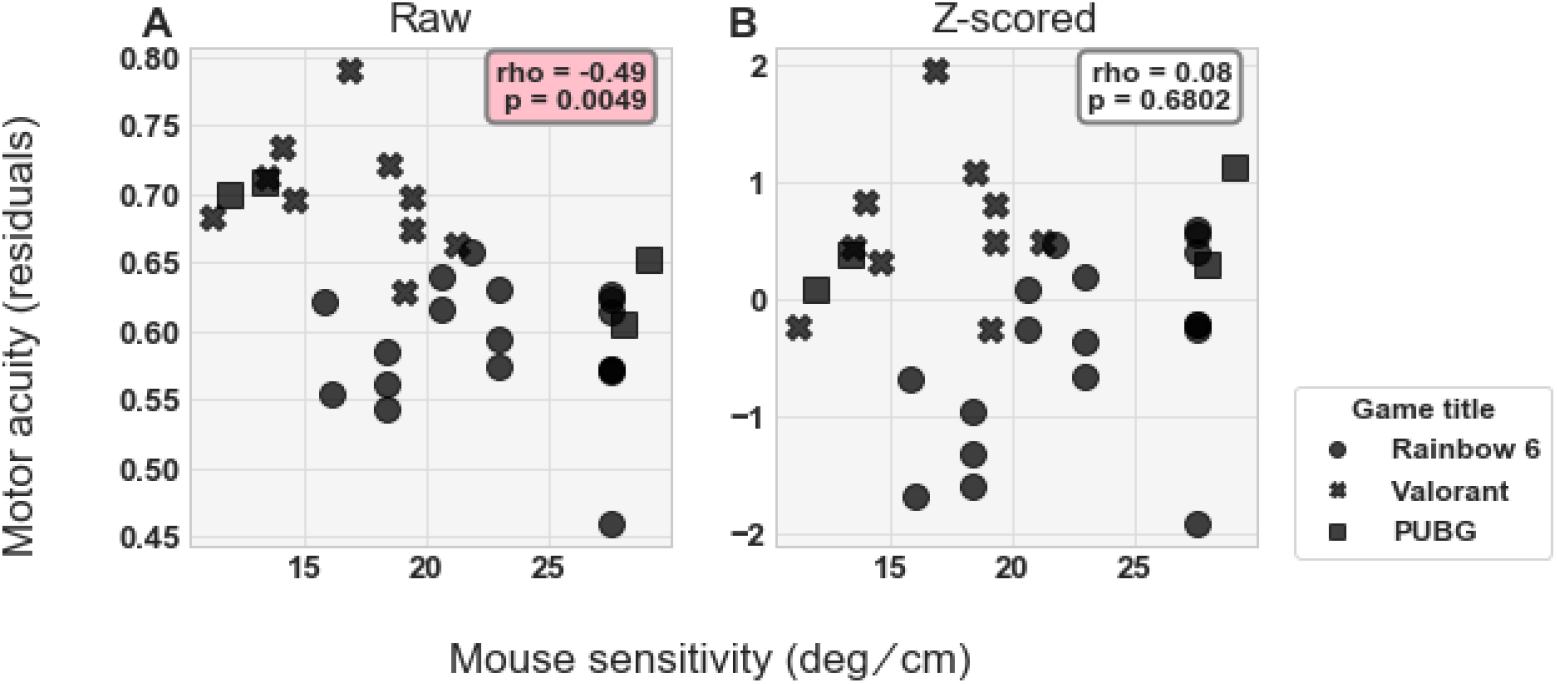
Correlation between motor acuity and mouse sensitivity: **A)** Before regressing out sensitivity, **B)** After regressing out sensitivity. Statistical results from Spearman’s Rank Correlation displayed in annotation boxes; where rho = Spearman’s Correlation Coefficient and p = *p*-value. Pink annotation boxes indicate a *p*-value *<* 0.05.

**Figure 10.**
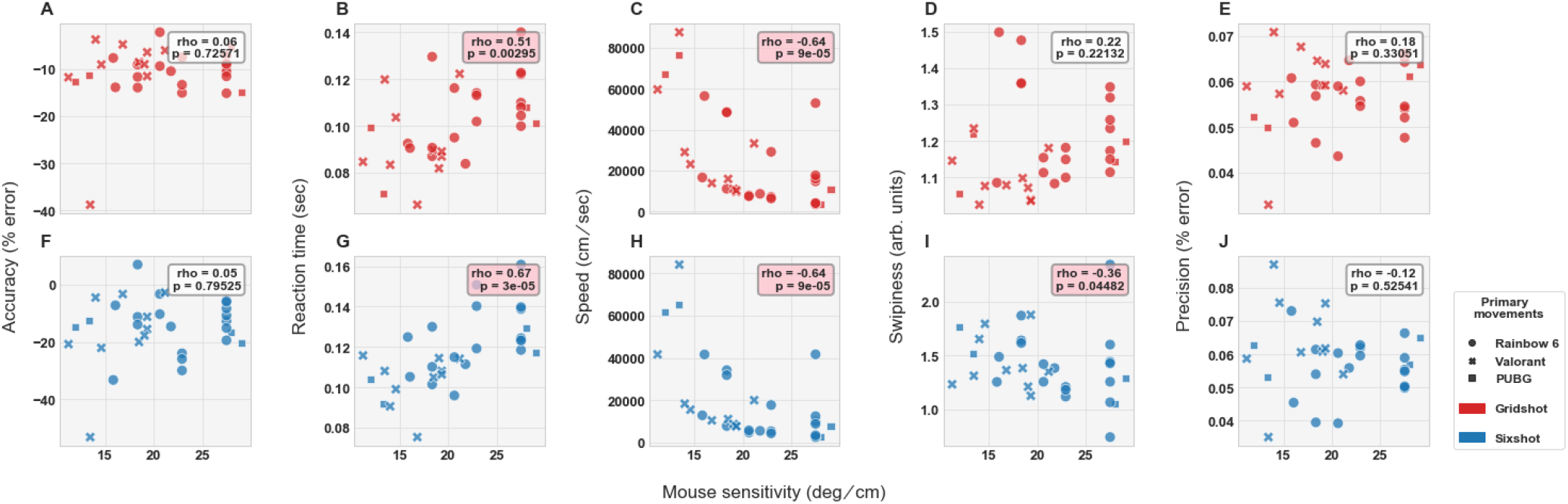
Correlation between movement kinematics (primary movements) and mouse sensitivity before regression for each metric in each task (see Legend): **A–E)** Gridshot, **F–J)** Sixshot. Participants marked by main game titled played (see Legend). Statistical results from Spearman’s Rank Correlation displayed in annotation boxes; where rho = Spearman’s Correlation Coefficient and p = *p*-value. Pink annotation boxes indicate a *p*-value *<* 0.05.

**Figure 11.**
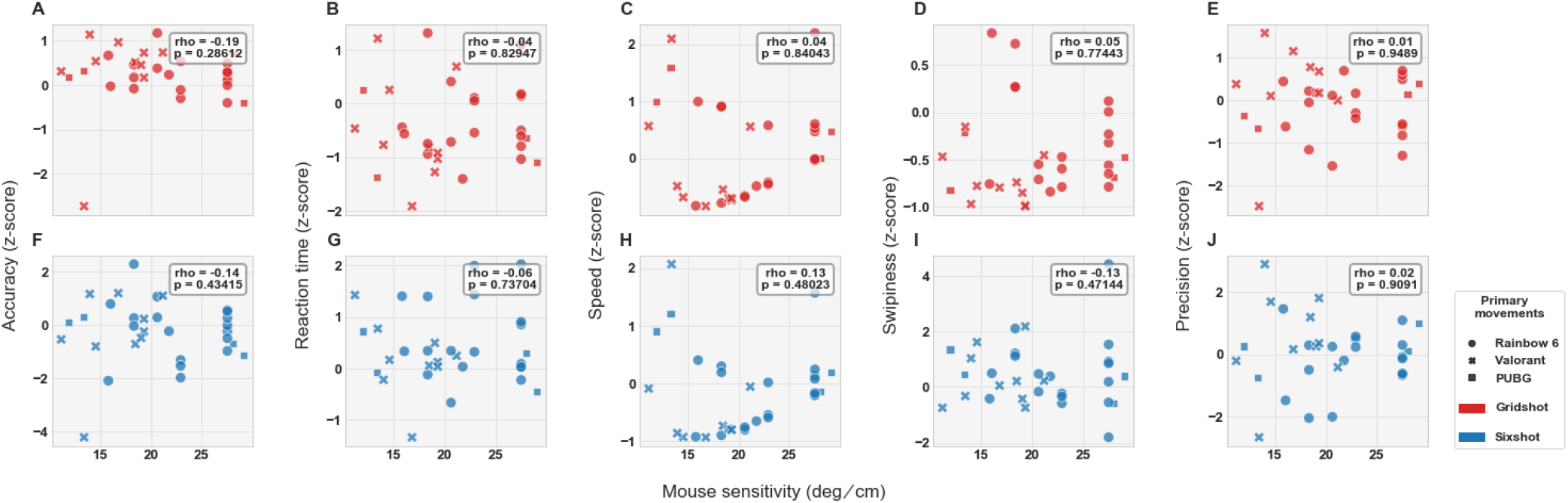
Correlation between movement kinematics (primary movements) and mouse sensitivity after regression for each metric in each task. Same format as Fig. 10.

**Figure 12.**
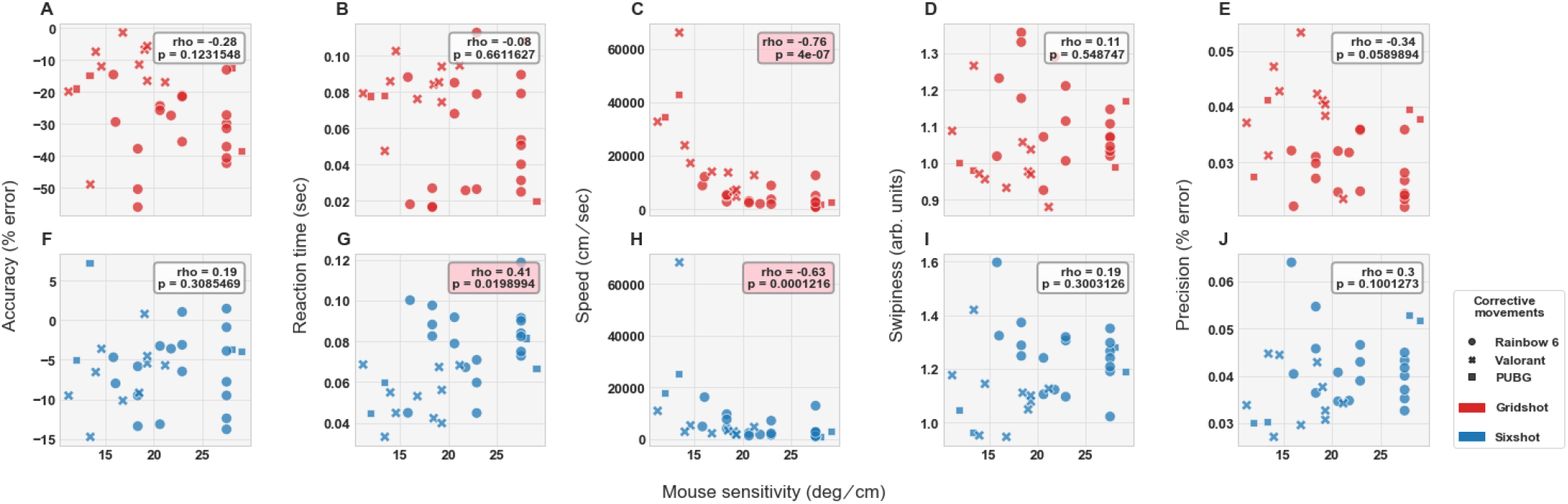
Correlation between movement kinematics (corrective movements) and mouse sensitivity before regression for each metric in each task. Same format as Fig. 10.

**Figure 13.**
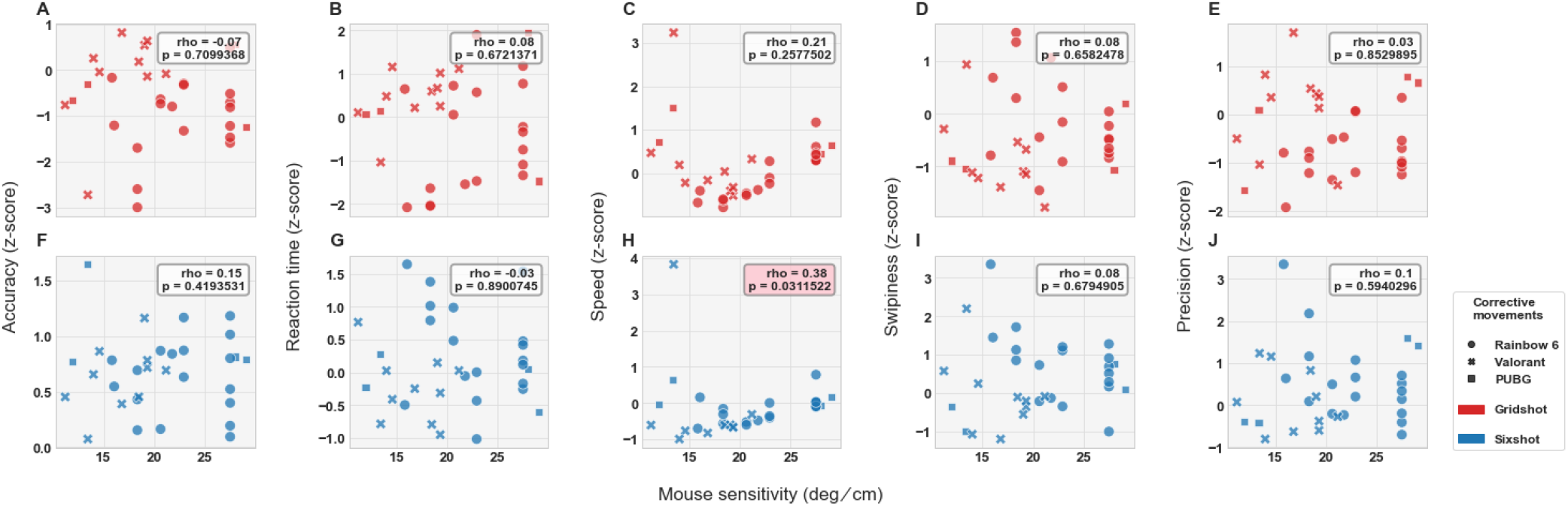
Correlation between movement kinematics (corrective movements) and mouse sensitivity after regression for each metric in each task. Same format as Fig. 10.

**Figure 14.**
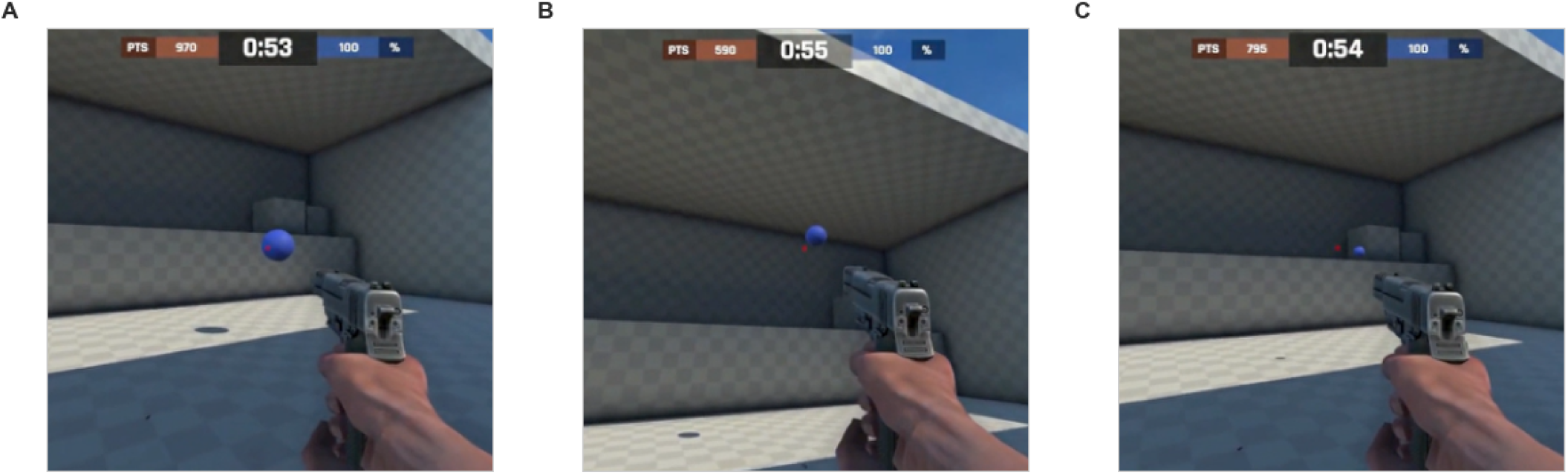
Sample screenshots of the Adaptive Reflexshot task. A) Large targets, B) Medium targets, and C) Small targets.

## REFERENCES

Adachi, P. J. and Willoughby, T. (2013a). Demolishing the competition: The longitudinal link between competitive video games, competitive gambling, and aggression. Journal of youth and adolescence 42, 1090–1104

Adachi, P. J. and Willoughby, T. (2013b). Do video games promote positive youth development? Journal of Adolescent Research 28, 155–165

Adachi, P. J. and Willoughby, T. (2013c). More than just fun and games: the longitudinal relationships between strategic video games, self-reported problem solving skills, and academic grades. Journal of youth and adolescence 42, 1041–1052

Bavelier, D., Green, C. S., Pouget, A., and Schrater, P. (2012). Brain plasticity through the life span: learning to learn and action video games. Annual review of neuroscience 35, 391–416

Brookes, J., Warburton, M., Alghadier, M., Mon-Williams, M., and Mushtaq, F. (2020). Studying human behavior with virtual reality: The unity experiment framework. Behavior research methods 52, 455–463

Burdea, G. C. (2003). Virtual rehabilitation–benefits and challenges. Methods of information in medicine 42, 519–523

Burke, J. W., McNeill, M., Charles, D., Morrow, P., Crosbie, J., and McDonough, S. (2009a). Serious games for upper limb rehabilitation following stroke. In 2009 Conference in Games and Virtual Worlds for Serious Applications (IEEE), 103–110

Burke, J. W., McNeill, M., Charles, D. K., Morrow, P. J., Crosbie, J. H., and McDonough, S. M. (2009b). Optimising engagement for stroke rehabilitation using serious games. The Visual Computer 25, 1085–1099

Campbell, M. J., Toth, A. J., Moran, A. P., Kowal, M., and Exton, C. (2018). esports: A new window on neurocognitive expertise? Progress in brain research 240, 161–174

Colzato, L. S., van den Wildenberg, W. P., Zmigrod, S., and Hommel, B. (2013). Action video gaming and cognitive control: playing first person shooter games is associated with improvement in working memory but not action inhibition. Psychological research 77, 234–239

Desmurget, M. and Grafton, S. (2000). Forward modeling allows feedback control for fast reaching movements. Trends in cognitive sciences 4, 423–431

Dye, M. W., Green, C. S., and Bavelier, D. (2009). Increasing speed of processing with action video games. Current directions in psychological science 18, 321–326

Egenfeldt-Nielsen, S., Smith, J. H., and Tosca, S. P. (2013). Understanding video games: The essential introduction (Routledge)

Fahle, M. (2005). Perceptual learning: specificity versus generalization. Current opinion in neurobiology 15, 154–160

Flatters, I., Hill, L. J., Williams, J. H., Barber, S. E., and Mon-Williams, M. (2014). Manual control age and sex differences in 4 to 11 year old children. PloS one 9, e88692

Funk, J. B. and Buchman, D. D. (1996). Playing violent video and computer games and adolescent self-concept. Journal of communication 46, 19–32

Gee, J. P. (2003). What video games have to teach us about learning and literacy. Computers in entertainment (CIE) 1, 20–20

Green, C. S. and Bavelier, D. (2003). Action video game modifies visual selective attention. Nature 423, 534–537

Green, C. S. and Bavelier, D. (2006). Enumeration versus multiple object tracking: The case of action video game players. Cognition 101, 217–245

Green, C. S. and Bavelier, D. (2007). Action-video-game experience alters the spatial resolution of vision. Psychological science 18, 88–94

Green, C. S. and Bavelier, D. (2008). Exercising your brain: a review of human brain plasticity and training-induced learning. Psychology and aging 23, 692

Heitz, R. P. (2014). The speed-accuracy tradeoff: history, physiology, methodology, and behavior. Frontiers in neuroscience 8, 150

Helgason, D., Francis, N., and Ante, J. (2005). Unity. Microsoft Windows [GAME ENGINE], Unity Technologies

Heuer, H. and Hegele, M. (2008). Constraints on visuo-motor adaptation depend on the type of visual feedback during practice. Experimental Brain Research 185, 101–110

Hong, J. S., Wasden, C., and Han, D. H. (2021). Introduction of digital therapeutics. Computer Methods and Programs in Biomedicine 209, 106319

Huang, J., Yan, E., Cheung, G., Nagappan, N., and Zimmermann, T. (2017). Master maker: Understanding gaming skill through practice and habit from gameplay behavior. Topics in cognitive science 9, 437–466

Ivory, J. D. (2015). A brief history of video games. The video game debate: Unravelling the physical, social, and psychological effects of digital games, 1–21

Jordan, M. I. and Rumelhart, D. E. (1992). Forward models: Supervised learning with a distal teacher. Cognitive science 16, 307–354

Kent, S. L. (2010). The Ultimate History of Video Games, Volume 1: From Pong to Pokemon and Beyond… the Story Behind the Craze That Touched Our Lives and Changed the World, vol. 1 (Crown)

Kowal, M., Toth, A. J., Exton, C., and Campbell, M. J. (2018). Different cognitive abilities displayed by action video gamers and non-gamers. Computers in Human Behavior 88, 255–262

Levenberg, K. (1944). A method for the solution of certain non-linear problems in least squares. Quarterly of applied mathematics 2, 164–168

Listman, J. B., Tsay, J. S., Kim, H. E., Mackey, W. E., and Heeger, D. J. (2021). Long-term motor learning in the “wild” with high volume video game data. Frontiers in human neuroscience 15

Maclean, N., Pound, P., Wolfe, C., and Rudd, A. (2000). Qualitative analysis of stroke patients’ motivation for rehabilitation. Bmj 321, 1051–1054

Maclean, N., Pound, P., Wolfe, C., and Rudd, A. (2002). The concept of patient motivation: a qualitative analysis of stroke professionals’ attitudes. Stroke 33, 444–448

Maniglia, M. and Seitz, A. R. (2018). Towards a whole brain model of perceptual learning. Current opinion in behavioral sciences 20, 47–55

Martin, T., Keating, J., Goodkin, H., Bastian, A., and Thach, W. (1996). Throwing while looking through prisms: I. focal olivocerebellar lesions impair adaptation. Brain 119, 1183–1198

McDougle, S. D. and Taylor, J. A. (2019). Dissociable cognitive strategies for sensorimotor learning. Nature communications 10, 1–13

Müller, H. and Sternad, D. (2004). Decomposition of variability in the execution of goal-oriented tasks: three components of skill improvement. Journal of Experimental Psychology: Human Perception and Performance 30, 212

Nudo, R. J. and Milliken, G. W. (1996). Reorganization of movement representations in primary motor cortex following focal ischemic infarcts in adult squirrel monkeys. Journal of neurophysiology 75, 2144–2149

Nudo, R. J., Milliken, G. W., Jenkins, W. M., and Merzenich, M. M. (1996a). Use-dependent alterations of movement representations in primary motor cortex of adult squirrel monkeys. Journal of Neuroscience 16, 785–807

Nudo, R. J., Wise, B. M., SiFuentes, F., and Milliken, G. W. (1996b). Neural substrates for the effects of rehabilitative training on motor recovery after ischemic infarct. Science 272, 1791–1794

Pedraza-Ramirez, I., Musculus, L., Raab, M., and Laborde, S. (2020). Setting the scientific stage for esports psychology: A systematic review. International Review of Sport and Exercise Psychology 13, 319–352

Plautz, E. J., Milliken, G. W., and Nudo, R. J. (2000). Effects of repetitive motor training on movement representations in adult squirrel monkeys: role of use versus learning. Neurobiology of learning and memory 74, 27–55

Pluss, M. A., Novak, A. R., Bennett, K., Panchuk, D., Coutts, A. J., and Fransen, J. (2020). Perceptual-motor abilities underlying expertise in esports. Journal of Expertise 3, 133–143

Popper, B. (2013). Field of streams: how twitch made video games a spectator sport. The Verge 30

Przybylski, A. K., Rigby, C. S., and Ryan, R. M. (2010). A motivational model of video game engagement. Review of general psychology 14, 154–166

PUBG Studios (2017). Pubg: Battlegrounds previously known as playerunknown’s battlegrounds. Microsoft Windows [GAME]

Putrino, D., Zanders, H., Rykman, A., Lee, P., Naeem, O., Disla, L., et al. (2015). Use of the gesaircraft video game for upper limb rehabilitation in stroke: A pilot study. Neurorehabil Neural Repair 30, 36–37

Riot Games (2020). Valorant. Microsoft Windows [GAME]

Shadmehr, R. and Mussa-Ivaldi, F. A. (1994). Adaptive representation of dynamics during learning of a motor task. Journal of neuroscience 14, 3208–3224

Shmuelof, L., Krakauer, J. W., and Mazzoni, P. (2012). How is a motor skill learned? change and invariance at the levels of task success and trajectory control. Journal of neurophysiology 108, 578–594

Shmuelof, L., Yang, J., Caffo, B., Mazzoni, P., and Krakauer, J. W. (2014). The neural correlates of learned motor acuity. Journal of neurophysiology 112, 971–980

Ubisoft (2015). Tom clancy’s rainbow six siege. Microsoft Windows [GAME]

Voida, A., Carpendale, S., and Greenberg, S. (2010). The individual and the group in console gaming. In Proceedings of the 2010 ACM conference on Computer supported cooperative work. 371–380

Wilterson, S. A. (2021). Implicit Recalibration and Strategic Compensation in Learned and De Novo Motor Skills. Ph.D. thesis, Princeton University

Winstein, C. J., Wolf, S. L., Dromerick, A. W., Lane, C. J., Nelsen, M. A., Lewthwaite, R., et al. (2016). Effect of a task-oriented rehabilitation program on upper extremity recovery following motor stroke: the icare randomized clinical trial. Jama 315, 571–581

Wolf, M. J. (2015). Video games around the world (MIT Press Cambridge, MA)

Wolf, S. L., Winstein, C. J., Miller, J. P., Taub, E., Uswatte, G., Morris, D., et al. (2006). Effect of constraint-induced movement therapy on upper extremity function 3 to 9 months after stroke: the excite randomized clinical trial. Jama 296, 2095–210

